# Neuroticism Heterogeneity Through Item-Level Associations in Resting-State Functional Connectivity

**DOI:** 10.1101/2025.05.23.655856

**Authors:** Masaya Misaki, Chun Chieh Fan, Wesley K Thompson, Heekyeong Park, Bohan Xu, Martin M Paulus

## Abstract

Neuroticism is characterized by emotional instability and increased susceptibility to stress-related disorders. While traditionally treated as a unitary construct, growing evidence suggests that neuroticism is heterogeneous in both its genetic basis and its effects on social and health outcomes. To quantify this heterogeneity at the neurofunctional level, we analyzed resting-state functional connectivity (RSFC) in a large sample (n = 33,180) from the UK Biobank dataset. Using machine learning regression analysis, we identified RSFC patterns associated with item-level responses from the neuroticism scale of the Eysenck Personality Questionnaire. The pattern of RSFC associations across questionnaire items reflected genetically defined clusters (Worry and Depressed Affect), showing a significant correlation (r = 0.767, p < 0.001). It also aligned with psycholometrically derived factors (Anxiety/Tension and Worry/Vulnerability), reflecting a factor structure consistent with prior psychological studies. These findings were replicated in a separate MRI scan session from the UK biobank dataset. Associated connectivity was primarily observed in cognitive control, sensory integration, and self-referential processing networks. These neurofunctional signatures position RSFC as a robust intermediate phenotype, bridging genetic predisposition for neuroticism and psychological states. This framework could enhance precision in predicting psychiatric vulnerability and inform tailored therapeutic interventions.

## Introduction

Neuroticism is a personality trait characterized by heightened negative affect, emotional instability, and susceptibility to stress ^1,2^. It is strongly associated with poor mental health outcomes ^3,4^, as individuals with high neuroticism face an increased risk of developing mood and anxiety disorders, underscoring its relevance in psychopathology ^5-9^.

While neuroticism is often treated as a unitary construct, evidence suggests it has a hierarchical substructure ^10^. Genetic analyses of item-level neuroticism responses suggest that neuroticism comprises multiple distinct genetic components, with some items more closely associated with depression and others with anxiety ^11,12^. Specifically, Nagel, et al. ^11^ identified two discrete clusters of neuroticism items in the Eysenck Personality Questionnaire ^13^ based on their genetic correlations in the UK Biobank ^14^. The ‘Depressed Affect’ cluster included the items *Loneliness/Isolation*, *Miserableness*, *Mood Swings*, and *Fed-Up Feelings*, while the ‘Worry’ cluster comprised *Nervous Feelings*, *Worry Too Long*, *Tense/Highly Strung*, and *Suffering from Nerves*.

Furthermore, bi-factor analyses have identified a general neuroticism factor alongside two specific factors, revealing that these components exert opposing effects on socioeconomic and health outcomes ^15,16^. Gale, et al. ^15^ and Hill, et al. ^16^ applied a bi-factor model analysis to neuroticism item responses in the UK Biobank ^14^, extracting continuous and overlapping factors rather than discrete clusters. They identified one general neuroticism factor and two specific factors: the ‘Anxiety/Tension’ factor, which had high loadings on *Nervous Feelings*, *Suffering from Nerves*, and *Tense/Highly Strung* items, and the ‘Worry/Vulnerability’ factor, which had high loadings on *Worry Too Long*, *Sensitivity/Hurt Feelings*, *Worrier/Anxious Feelings*, and *Guilty Feelings* items. Hill, et al. ^16^ further identified genetic associations of these factors. Interestingly, the general and specific factors had opposite influences on health and socioeconomic status. While the general factor was associated with adverse outcomes, the specific factors were linked to high socioeconomic status, high intelligence, self-rated good health, and increased longevity. These findings emphasize the need to investigate neuroticism as a multifaceted trait, with each facet examined separately to determine its contribution to health outcomes.

Given the heterogeneity of neuroticism and its distinct associations with social and health outcomes, including depression and worry, further research is needed to clarify how its components are reflected in brain function and contribute to individual differences ^17^. This will help elucidate the mechanistic links between personality traits and cognitive brain processes, with important implications for understanding vulnerability to mood and anxiety disorders, which are associated with high levels of neuroticism ^5,18^. One promising avenue for understanding the neurofunctional basis of neuroticism is resting-state functional connectivity (RSFC), which examines intrinsic brain network organization and functional interactions between brain regions in the absence of external tasks. Prior research indicates that neuroticism is associated with widespread alterations in RSFC. Notably, differences in connectivity between the amygdala and regions such as the precuneus and insula may reflect differences in socio-emotional processing and stress reactivity among individuals with high neuroticism ^19-22^. Network analyses using graph theory have highlighted that highly neurotic individuals exhibit a functional network architecture that deviates from the typical small-world organization, instead displaying more random connectivity patterns ^23^. These differences are particularly pronounced in networks involved in emotion regulation and salience detection, suggesting a neurobiological framework through which neuroticism influences emotional and cognitive functioning. Furthermore, connectome-based predictive modeling (CPM) has demonstrated that neuroticism can be predicted based on RSFC patterns ^24^, highlighting the utility of functional connectivity analyses in capturing trait-level personality differences.

Despite these advances, most prior studies have treated neuroticism as a single trait, typically deriving a composite score by summing responses across all items on a neuroticism scale. While high neuroticism is a well-established risk factor for both depressive and anxiety disorders ^5,18^, specific subcomponents of neuroticism may have distinct associations with these conditions, a distinction supported by genetic research ^11,16^.

Although these studies suggest that neuroticism consists of multiple biologically distinct subcomponents, significant gaps remain in our understanding of how this heterogeneity interacts with neurofunctional architecture. While Nagel, et al. ^12^ investigated the neural associations of these genetic patterns at the cell level, the functional organization of the brain related to neuroticism subcomponents has not been examined at the whole-brain network level. To address this gap, the present study investigated RSFC associations with item-level responses from the neuroticism questionnaire in the UK Biobank and examined the relationships between items to determine whether similarly distinct clusters or overlapping factors were observed in their RSFC associations.

We employed a predictive modeling approach ^25^, where a machine learning regression analysis was used to predict questionnaire scores from RSFC patterns, to examine whole-brain RSFC associations with individual responses to each neuroticism questionnaire item. While Hsu, et al. ^24^ previously applied a similar approach to the overall neuroticism score, we conducted this analysis at the item level. Next, we identified clusters of items based on similarities in their RSFC associations, following the approach of Nagel, et al. ^11^ for genetic associations. Additionally, we applied a bi-factor model analysis ^26,27^ to the RSFC patterns associated with each item to extract both general and specific factors, following the approach of Gale, et al. ^15^ and Hill, et al. ^16^. Finally, we identified the RSFC patterns associated with these clusters and factors to elucidate their neurofunctional architecture.

## Methods

### Samples

This study analyzed data from participants in the UK Biobank study (http://www.ukbiobank.ac.uk) ^14^ under data application 95683. fMRI data were obtained from the first imaging instance of 33,180 participants (17,269 females) between the ages of 44 and 82 years (mean = 63.7, SD = 7.7), all of whom had completed the neuroticism questionnaire. Data from the second imaging instance, consisting of 3,748 participants (1,919 females) aged 49 to 83 years (mean = 64.0, SD = 7.2), were used to assess the replicability and stability of the results.

### Resting-state functional connectivity (RSFC)

RSFC analysis was performed using resting-state fMRI (rfMRI) partial correlation matrices derived from a 100-dimensional group independent component analysis (ICA) decomposition. Data were generated from preprocessed rfMRI images (TR/TE = 735/39 ms, voxel size = 2.4 mm³, matrix = 88 × 88 × 64, duration = 6 min 10 s). During preprocessing, ICA was applied to individual samples, and noise components were identified and removed using FMRIB’s ICA-based X-noiseifier (FIX). The preprocessed images were normalized to the MNI template brain and group ICA was then performed to extract 100 components, which were classified into signal and noise components. The time courses of these signal components were computed using spatial regression of the component maps, resulting in the extraction of 55 component signals. Finally, partial correlations between these signals were calculated as RSFC measures and normalized by the average correlation of random noise signals with similar autocorrelation properties. For further details, see Alfaro-Almagro, et al. ^28^.

### Neuroticism scores

Neuroticism was assessed using the Eysenck Personality Questionnaire (12 items) ^13^. Each item had four possible response options: “Yes,” “No,” “Do not know,” and “Prefer not to answer.” Only participants who responded “Yes” or “No” to all 12 questions were included in the analyses. *Predictive modeling of neuroticism item responses using RSFC* A machine learning (ML) predictive modeling approach ^25^ was employed to examine the association between RSFC and neuroticism item responses. Specifically, a classification analysis with RSFC was conducted on yes/no responses for each neuroticism questionnaire item. Model training and evaluation were performed using three-fold cross-validation, where the dataset was split into three groups: two groups were used for model optimization, while the remaining hold-out group was used for evaluation.

To control for potential confounding factors, the covariates of scan location, sex, age, squared age, mean motion, signal-to-noise ratio, and T1-EPI discrepancies were regressed out from the RSFC values using a linear model. Scan location and sex effects were first regressed out from each RSFC value, followed by the regression of the remaining covariates within each scan location and sex allowing covariate effects to differ across groups ^29^. Covariate regression was performed only within the training data, and the fitted regression model was applied to the test data to prevent information leakage ^30^. The residuals from this regression were used as inputs for the ML analysis.

For ML classification, we utilized an automated machine learning (AutoML) approach. AutoML addresses the Combined Algorithm Selection and Hyperparameter (CASH) optimization problem by automatically optimizing feature selection, model algorithm, and hyperparameters. Specifically, we used the H2O AutoML software package ^31^ (https://docs.h2o.ai/h2o/latest-stable/h2o-docs/automl.html), which is known for its comprehensive algorithm selection and optimization capabilities.

H2O AutoML evaluated various models, including Generalized Linear Models (GLM), Gradient Boosting Machines, Random Forest, Naïve Bayes, Support Vector Machine, and Deep Neural Networks, as well as their ensembles. Parameter optimization was performed using nested cross-validation within the training data, where the training set was further split into three sub-groups. The final model was then trained on the entire training dataset using the optimized CASH parameters.

### Model evaluations

Model performance was evaluated using the Area Under the Receiver Operating Characteristic Curve (AUC) to mitigate bias caused by imbalanced yes/no response ratios across items. The AUC value was computed across the test samples using multiple cross-validation repetitions. The statistical significance of the AUC values was assessed using a normal approximation of the Mann-Whitney U statistic ^32^.

The contribution of each functional connectivity (FC) feature to predicting neuroticism item response was evaluated by measuring feature importance, averaged across cross-validation folds. Since the regularized logistic regression model (GLM) was consistently selected as the best-performing model, feature importance was assessed based on standardized coefficients.

### Comparison of RSFC Associations with Genetic Associations and Bi-Factor Structures of Neuroticism

We compared the similarity structures between items, one derived from genetic correlations and the other from RSFC associations. RSFC similarity between items was defined as the Pearson correlation between the standardized regression coefficients for each item response, while genetic correlations were obtained from Nagel, et al. ^11^. We then calculated the Pearson correlation between the RSFC-based and genetically derived similarities between items.

To further explore the underlying neurofunctional structure, we applied an exploratory bi-factor model analysis ^26,27^ to the RSFC associations of individual items. The input to the factor analysis was a matrix of items by standardized regression coefficients derived from RSFC-based regression models. Following the approach of Gale, et al. ^15^ and Hill, et al. ^16^, the bi-factor analysis extracted one general factor and two specific factors. The analysis was conducted using the ‘psych’ ^33^ and ‘GPArotation’ ^34^ packages in R ^35^. Factor extraction was performed using the principal axis method with ‘bifactorq’ oblique rotation.

### Assessment of replicability and stability of results

To assess the replicability of RSFC association structures for neuroticism item responses, we assessed whether the model trained on the first MRI instances could generalize to the second. Participants from the second instance were assigned to the same cross-validation scheme used in training with the first instance. To prevent information leakage, we ensured that each test used a model that had not been trained on the same participants’ data from the other instance. We also performed RSFC predictive modeling on data from the second MRI instance, following the same procedure as in the first instance. We then examined the RSFC associations in relation to genetic associations and factor structures.

The stability of neuroticism responses across instances was assessed at the individual participant level for both questionnaire item responses and RSFC associations. Stability was evaluated for sum scores of the Worry and Depressive Affect clusters ^11^ and factor scores derived from an exploratory bi-factor analysis. For item responses, binary values (yes = 1, no = 0) were used to calculate the sum scores of clustered items and the factor scores for the general and two specific factors. Factor loadings were taken from Hill, et al. ^16^, and factor scores were computed using the regression (Thurstone’s) method.

For RSFC associations, individual participant responses were calculated using the ML model output of class probabilities, which were transformed into z-values via logit transformation. The model trained on the first MRI instance was used for this calculation. These values were then used to compute cluster sum and factor scores, with factor loadings derived from the exploratory bi-factor analysis of RSFC associations.

The agreement between instances was evaluated using the intraclass correlation coefficient (ICC) with a two-way random-effects model, using the ‘irr’ package ^36^ in R ^35^.

## Results

### Classification of Neuroticism Item Responses from RSFC

The AutoML analysis for classifying neuroticism item responses from RSFC selected the regularized logistic regression model in GLM as the best-performing model across all items and cross-validation folds. Figure 1 presents the AUC values for classification performance (left) along with their *p*-values (right). While AUC values were only slightly above 0.5, their *p*-values remained highly significant, even under a strict threshold (*p* < 5e^-8^), except for the *irritability* item (*p* = 7e^-5^).

**Figure 1.**
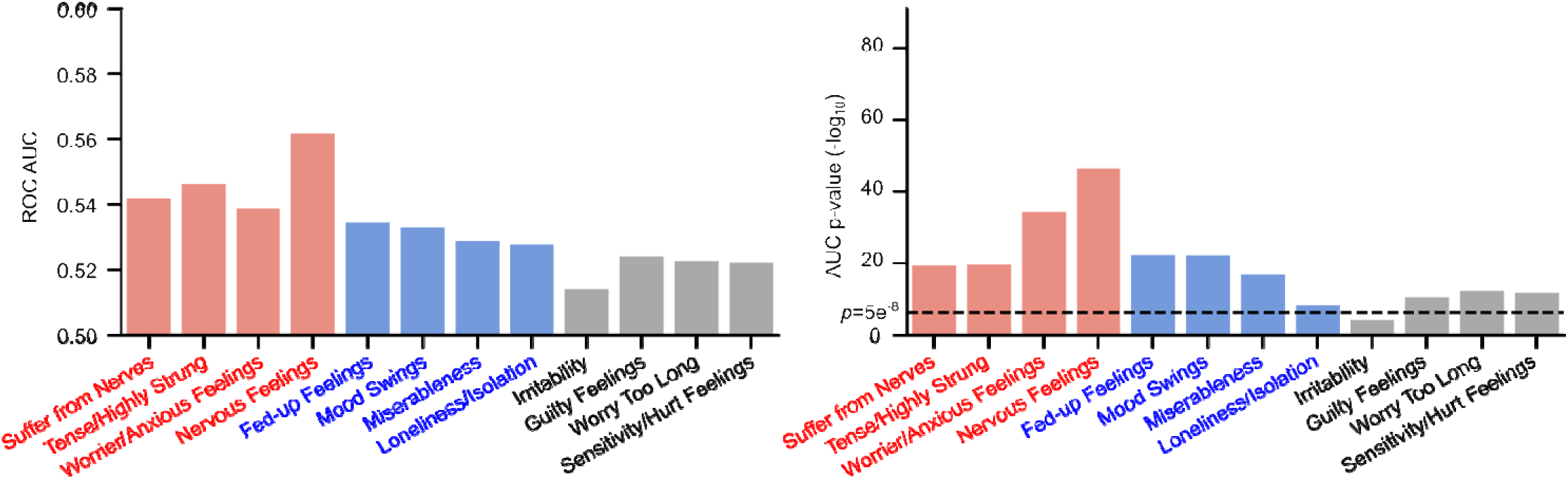
Classification performance (Area Under the Receiver Operating Characteristic Curve; AUC) and corresponding *p*-values for neuroticism item responses using AutoML. Colors indicate item clusters identified based on their genetic correlations ^11^: Red represents the Worry cluster, while Blue represents the Depressed Affect cluster.

### Alignment of genetic and RSFC-based item clusters

Figure 2 presents the correlation matrix of RSFC-based regression coefficient patterns across items. Correlations were computed between the standardized coefficients of the logistic regression models for each item. The matrix revealed clustered patterns corresponding to the Worry and Depressed Affect clusters, consistent with genetic correlation patterns reported by Nagel, et al. ^11^.

**Figure 2.**
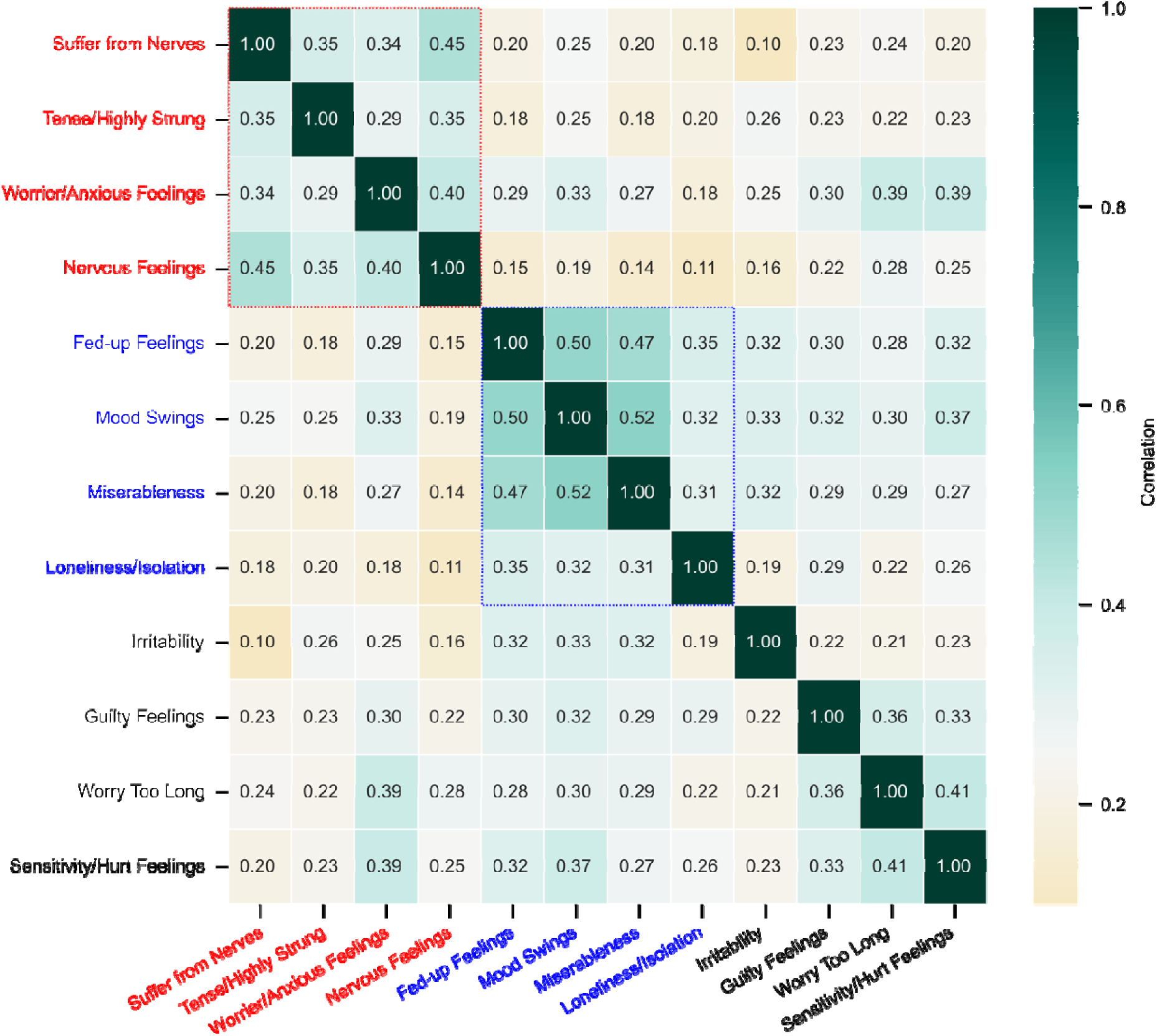
Correlation matrix of resting-state functional connectivity (RSFC) associations between neuroticism items. The colors of the row and column labels indicate item clusters identified based on their genetic correlations ^11^: Red represents the Worry cluster, while Blue represents the Depressed Affect cluster.

The alignment of RSFC and genetic associations was further confirmed by their correlation strength. Figure 3 illustrates the relationship between RSFC pattern correlations across items and their genetic correlations obtained from Nagel, et al. ^11^. The RSFC-based item similarity structure was highly correlated with the genetic correlation structure (*r* = 0.767, *p* < 0.001), suggesting that neurofunctional associations in RSFC capture the genetic heterogeneity underlying neuroticism.

**Figure 3.**
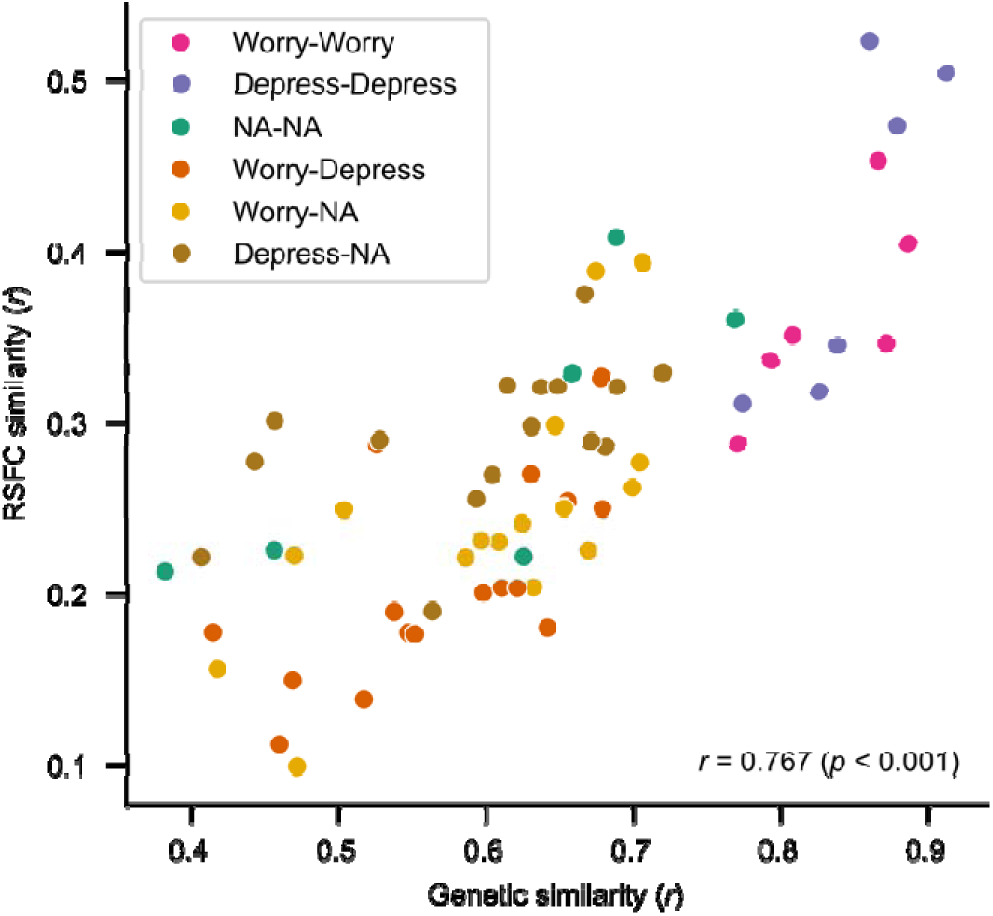
The relationship between resting-state functional connectivity (RSFC) pattern correlations across items and their genetic correlations obtained from Nagel, et al. ^11^. Dot colors indicate the combinations of item clusters, while NA denotes items that were not assigned to either cluster.

### Bi-factor analysis of RSFC associations

The bi-factor analysis extracted one general factor and two specific factors from the RSFC association patterns of the items. These factors explained 29%, 7%, and 3% of the variance, respectively. Model fit indices indicated good fit, with a root mean square of the residuals (RMSR) of 0.02 and a Tucker–Lewis Index (TLI) of 0.961.

Figure 4 presents the factor loading patterns. The first specific factor (F1) showed higher loadings on *Nervous Feelings*, *Suffer from Nerves*, *Tense/Highly Strung*, and *Worrier/Anxious Feelings* items, while the second specific factor (F2) had higher loadings on *Worry Too Long, Sensitivity/Hurt Feelings*, and *Guilty Feelings*. These factor structures closely aligned with the Anxiety/Tension (F1) and Worry/Vulnerability (F2) factors identified in previous studies ^15,16^, except for *Worrier/Anxious Feelings*, which previously loaded onto the Worry/Vulnerability factor (F2) in questionnaire-based models ^15,16^.

**Figure 4.**
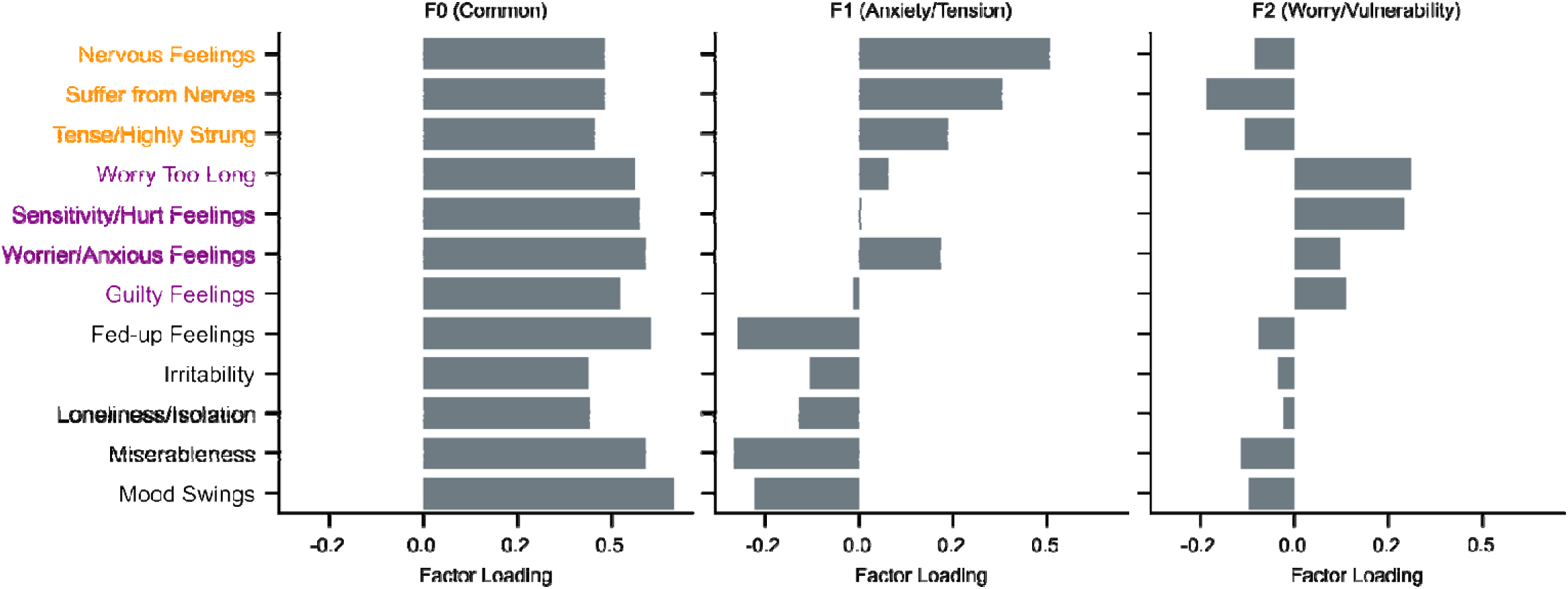
Factor loadings for the general factor (F0) and two specific factors (F1, F2) extracted through exploratory bi-factor analysis. The colors of the row item labels indicate the specific factors identified based on questionnaire responses by Hill, et al. ^16^: Orange represents the Anxiety/Tension factor, while Purple represents the Worry/Vulnerability factor.

### RSFC patterns associated with the neuroticism item clusters

Figure 5 presents the representative FCs associated with the Worry and Depressed Affect clusters of neuroticism items. RSFC was calculated between independent component (IC) signal time series, with each node representing an IC. Connection values were derived from the average z-values of the GLM coefficients across questionnaire items within each cluster. The plot visualizes only connections with an absolute z-value greater than 2.576 (*p* < 0.01), excluding FCs assigned a weight of 0 in any cross-validation fold due to regularization. This selection enhances presentation clarity by highlighting the most representative FCs.

**Figure 5.**
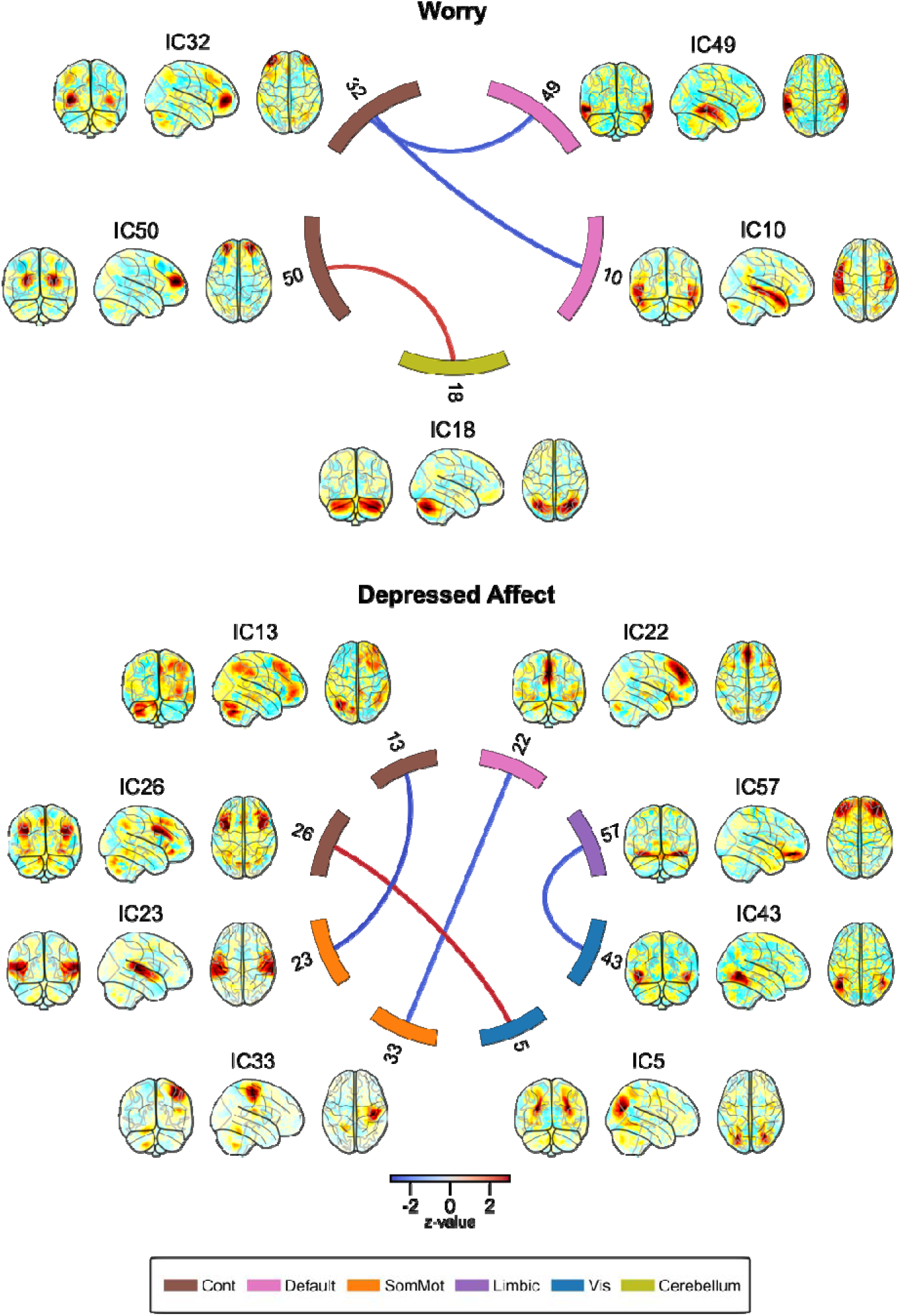
Representative resting-state functional connectivity associated with the Worry and Depressed Affect clusters of neuroticism items. The plot visualizes only connections with an absolute z-value greater than 2.576 (p < 0.01). The IC weight map for each node is shown on the glass brain. Node numbers indicate the index of the independent component (IC), and node colors represent network labels. The network labels were based on Yeo, et al. ^37^: Cont, Frontoparietal Control network; Default, Default mode network; SomMot, Somatomotor network; Limbic, Limbic network; Vis, Visual network; Cerebellum, Cerebellum network.

While Figure 5 focuses on key FCs, ML-based prediction with regularized logistic regression relies on a broader network of FCs, and a more comprehensive view of RSFC patterns is provided in Supplementary Figure S1. The independent components corresponding to FC nodes were assigned to resting-state networks based on Yeo, et al. ^37^ and were manually categorized into subcortical and cerebellar regions.

For the Worry cluster, representative RSFCs included reduced FC between the IC of the prefrontal regions in the control network (IC32) and ICs of the middle temporal regions (IC49) and the middle temporal and hippocampal regions (IC10), which are associated with episodic memory retrieval in the default mode network. Additionally, increased FC was observed between the ICs of the superior frontal regions in the control network (IC50) and the cerebellum (IC18).

For the Depressed Affect cluster, reduced FC was observed between the ICs in the parieto-frontal control regions of the control network (IC13) and the superior temporal regions of the somatomotor network (IC23), which are associated with auditory processing; between the ICs in the medial superior frontal regions of the default mode network (IC22) and the right primary somatosensory/motor region in the somatomotor network (IC33); and between the ICs in the orbitofrontal regions of the limbic network (IC57), often linked to emotion processing, and the lateral inferior occipital to middle temporal regions of the visual network (IC43). Additionally, increased FC was observed between the ICs in the middle frontal regions of the control network (IC26) and the lateral middle occipital regions of the visual network (IC5).

### RSFC patterns associated with neuroticism factors

Figure 6 presents the representative FCs associated with the factors of RSFC associations for neuroticism items, extracted through exploratory bi-factor analysis. Connection values were calculated as the weighted sum of the z-values of the GLM coefficients, using each factor’s weight for each item. To enhance clarity and highlight representative FCs, the plot visualizes only the top four connections with the highest absolute weighted z-values. A more comprehensive view of the RSFC patterns is provided in Supplementary Figure S2.

**Figure 6.**
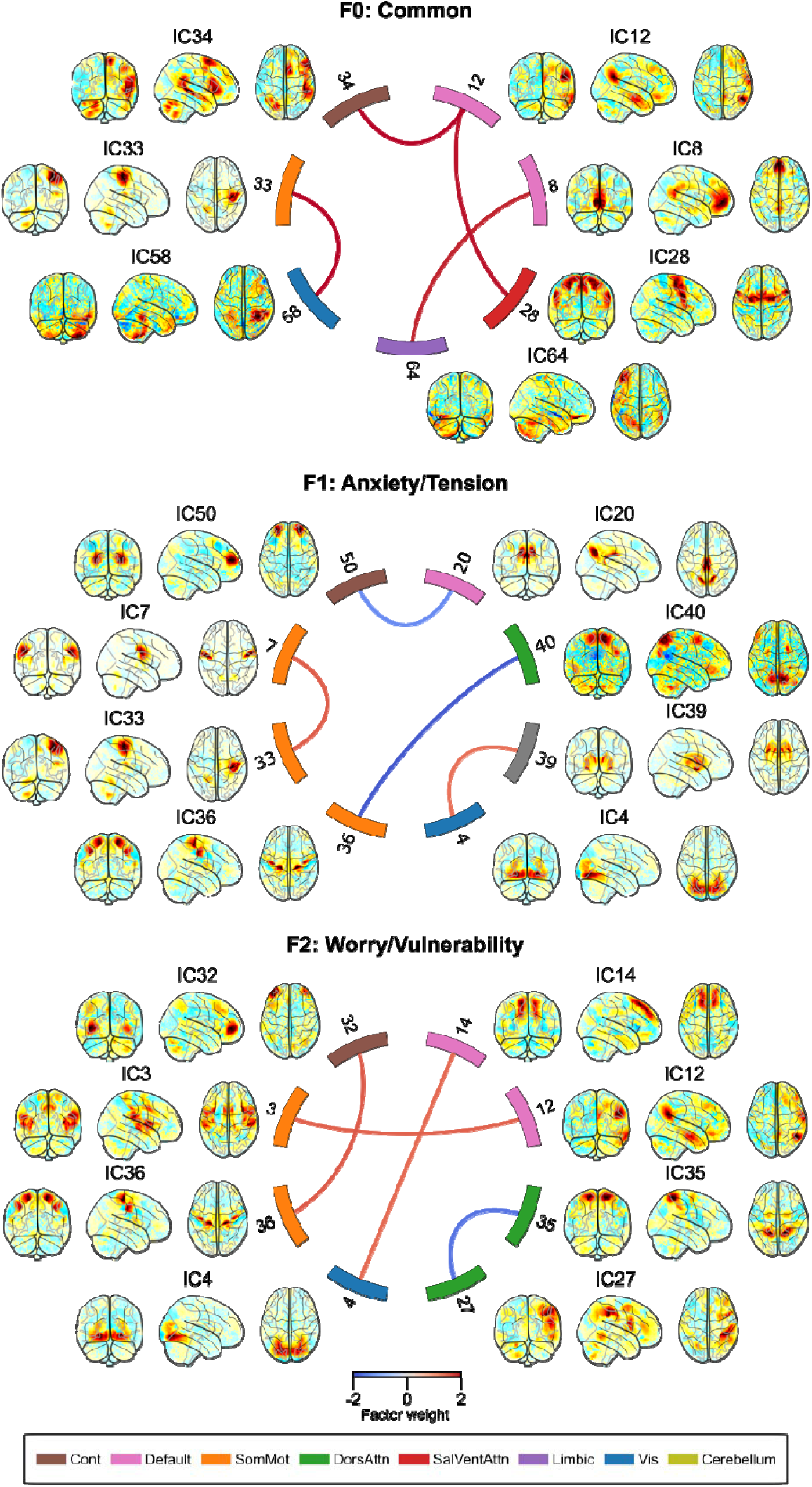
Representative resting-state functional connectivity (RSFC) associated with the factors of RSFC associations for neuroticism items, extracted through exploratory bi-factor analysis. The plot visualizes only the top four connections with the highest absolute weighted z-values. The IC weight map for each node is shown on the glass brain. Node numbers indicate the index of the IC, and node colors represent network labels. The network labels were based on Yeo, et al. ^37^: Cont, Frontoparietal Control network; Default, Default mode network; SomMot, Somatomotor network; DorsAttn, Dorsal attention network; Sal/VentAttn, Salience/Ventral attention network; Limbic, Limbic network; Vis, Visual network; Cerebellum, Cerebellum network.

For the general factor (F0), the associated FCs included increased connectivity between the right temporoparietal junction (TPJ)/middle temporal region in the default mode network (IC12) and the right superior frontal, ventrolateral prefrontal, pre-SMA, and STS regions in the control network (IC34), as well as the lateral superior premotor regions and supplementary motor area (SMA) in the salience network (IC28); between the core regions of the default mode network, including the medial prefrontal cortex, precuneus, and inferior parietal regions in the default mode network (IC8), and the left orbitofrontal region (IC64); and between the right primary somatosensory/motor region in the somatomotor network (IC33) and the right inferior temporal region in the visual network (IC58).

For the first specific factor (F1), corresponding to Anxiety/Tension, the associated FCs included decreased connectivity between the posterior cingulate cortex (PCC) and the precuneus in the default mode network (IC20) and the anterior superior prefrontal regions in the control network (IC50); between the primary somatosensory motor cortex in the somatomotor network (IC36) and the superior parietal regions in the dorsal attention network (IC40); as well as increased connectivity between the bilateral primary somatosensory/motor region in the somatomotor network (IC7) and the right primary somatosensory/motor region in the somatomotor network (IC33); and between the basal ganglia regions in the subcortical area related to motor and emotion processing (IC39) and the lingual/fusiform gyrus in the visual network (IC4).

For the second specific factor (F2), corresponding to Worry/Vulnerability, the associated FCs included increased connectivity between the prefrontal regions in the control network (IC32) and the primary somatosensory motor cortex in the somatomotor network (IC36); between the superior frontal regions in the default mode network (IC14) and the lingual/fusiform gyrus in the visual network (IC4); and between the right TPJ/middle temporal region in the default mode network (IC12) and the insula regions in the somatomotor network (IC3). Decreased connectivity between the superior parietal regions in the dorsal attention network (IC35) and the right inferior frontal to intraparietal sulcus regions in the dorsal attention network (IC27) was also observed.

### Replicability of the RSFC association structure and stability of neuroticism responses

The ML model trained on data from the first MRI instance generalized to the second MRI instance. Supplementary Figure S3 shows the AUC of models trained on the first MRI instance and applied to the second MRI instance. The model performed similarly across the first (Fig. 1) and second MRI instances.

The alignment between RSFC association and genetic correlation structures was replicated in the predictive modeling analysis using data from another MRI instance. The RSFC-based item similarity structure remained highly correlated with the genetic correlation structure (*r* = 0.750, *p* < 0.001, Supplementary Figure S4). Factor loadings also showed a consistent structure for the specific factors (Supplementary Figure S5), with the largest item loadings aligning with the Anxiety/Tension and Worry/Vulnerability factor structures, except for the Worrier/Anxious Feelings item, which showed the same pattern as in the first MRI instance (Fig. 4). These findings indicate that RSFC associations with neuroticism item responses were consistent across MRI instances.

Regarding the stability of questionnaire responses at the individual participant level, the Worry and Depressed Affect cluster scores demonstrated good agreement across sessions, with ICC = 0.792 (95% confidence interval [CI] = 0.779–0.805) for the Worry cluster and ICC = 0.743 (CI = 0.727–0.758) for the Depressed Affect cluster. The general factor score also showed good stability (ICC = 0.790, CI = 0.776–0.802), while the specific factor scores were moderately stable, with ICC = 0.641 (CI = 0.620–0.661) for the Anxiety/Tension factor and ICC = 0.671 (CI = 0.652–0.690) for the Worry/Vulnerability factor.

In contrast, the stability of individual prediction scores from the RSFC-based regression model across sessions was poor. The ICC values were 0.385 (CI = 0.335–0.414) for the Worry cluster, 0.262 (CI = 0.229–0.294) for the Depressed Affect cluster, 0.354 (CI = 0.323–0.384) for the general factor, 0.330 (CI = 0.299–0.361) for the Anxiety/Tension factor, and 0.223 (CI = 0.190–0.256) for the Worry/Vulnerability factor.

## Discussion

This study investigated how the neurofunctional heterogeneity of neuroticism is reflected in RSFC, particularly in relation to distinct subcomponents identified through genetic and psychometric analyses. The findings indicate that neuroticism is associated with distinct yet overlapping RSFC patterns that closely align with genetically informed clusters (e.g., Worry and Depressed Affect) and psychometrically derived factors (e.g., Anxiety/Tension and Worry/Vulnerability). These patterns involve differential connectivity in networks related to cognitive control, sensory integration, and self-referential processing. The results suggest that RSFC serves as an intermediate phenotype, capturing both heritable traits and psychological states associated with neuroticism.

The most parsimonious mechanistic explanation emerging from this study is that neuroticism, as a multifaceted personality trait, can be at least partially instantiated via distinct but interrelated neurofunctional connectivity patterns that correspond to both genetically informed subcomponents and self-reported psychological experiences. RSFC analyses revealed that neurofunctional associations of neuroticism item responses closely align with previously identified genetic clusters (e.g., Worry and Depressed Affect) and psychometrically derived factors (e.g., Anxiety/Tension and Worry/Vulnerability). These results reinforce the idea that neuroticism’s heterogeneity is reflected at the neural level, bridging genetic predispositions with cognitive-emotional processes.

The alignment of RSFC with these subcomponents suggests that different aspects of neuroticism are associated with distinct variations in functional networks. The Worry cluster appears to be linked to decreased coordination between self-referential and cognitive control systems, as evidenced by reduced connectivity between prefrontal control regions and DMN-related areas. This reduced connectivity may reflect differences in top-down modulation of memory retrieval and internally directed thought, which could relate to tendencies toward rumination ^38^. Additionally, increased connectivity between executive control regions in the superior frontal cortex and the cerebellum suggests variability in cognitive regulation ^39-42^. This pattern may reflect compensatory engagement in response to worry-related demands or less efficient regulation contributing to sustained cognitive engagement with negative content. Notably, differences in cerebellar FC have been observed in individuals with major depressive disorder (MDD) ^43^ and have been associated with repetitive negative thinking ^44^, suggesting that cerebellar connectivity patterns may also be relevant to worry-prone cognitive style.

In contrast, the Depressed Affect cluster involves distinct connectivity patterns in control, limbic, and sensory-motor networks, pointing to differences in sensory integration, emotion processing, and motor regulation. Reduced connectivity between orbitofrontal regions and sensory regions (visual and auditory) may reflect altered integration of affective and perceptual information, potentially related to slowed emotional and behavioral responses often observed in mood disorders ^45,46^.

The bi-factor analysis further clarifies these associations by revealing distinct RSFC patterns corresponding to general and specific neuroticism components. The general factor (F0) was associated with increased connectivity among the default mode, control, and salience networks, suggesting broader involvement in self-referential processing, executive control, and attention modulation ^47,48^. Additionally, elevated somatomotor-visual connectivity may correspond to enhanced perceptual processing or vigilance, which are characteristics sometimes associated with high negative affect ^49,50^.

The Anxiety/Tension factor (F1) was characterized by reduced connectivity between the DMN and prefrontal control networks, and between sensorimotor and attentional networks, suggesting variation in self-referential and sensory-attentional coordination ^51-53^. Enhanced connectivity between the basal ganglia and visual areas may reflect heightened vigilance or increased sensitivity to environmental stimuli, which are consistent with anxiety-prone cognitive patterns ^54,55^.

The Worry/Vulnerability factor (F2) showed increased connectivity between prefrontal control and sensorimotor regions, suggesting enhanced regulatory engagement in motor domains, possibly linked to behavioral avoidance tendencies. Stronger superior frontal–visual connectivity may indicate increased perceptual responsiveness to emotionally salient stimuli, reinforcing sustained attentional focus on potential threats ^49,50^. Elevated TPJ–insula connectivity may support increased interoceptive awareness and social-cognitive processing associated with worry and sensitivity to environmental uncertainty ^56,57^, while reduced dorsal attention network connectivity may suggest challenges in attentional disengagement, a trait commonly associated with high worry ^58,59^.

Taken together, these RSFC patterns illustrate that neuroticism is not a unitary construct but is composed of diverse neural signatures that correspond to its various behavioral, cognitive, and affective dimensions.

These cluster and factor structures were generalized and replicated in MRI data from a separate imaging session, confirming the robustness of the findings at the group level. However, at the individual level, the prediction scores derived from the RSFC-based model showed low stability across sessions. Given the high stability of questionnaire responses, this variability is unlikely to reflect changes in the personality trait itself. Rather, it suggests that RSFC is sensitive to momentary psychological states, which may obscure stable trait-level characteristics. Nonetheless, the machine learning model generalized well, indicating that it likely captured trait-related variance in RSFC. However, the relatively low predictive performance for neuroticism item responses, although statistically significant, suggests that the trait-related component of RSFC variance is modest.

Prior research has similarly shown that RSFC’s predictive power for individual traits is generally low ^60^, particularly in short-term recordings, and that scans lasting several hours or more are required to obtain stable individual-specific patterns ^61^. While this study demonstrates that multivariate pattern analysis using ML can extract group-level relationships from RSFC data, its applicability to individual-level prediction remains uncertain. Future research may need to incorporate prolonged resting-state recordings to improve the reliability of RSFC-based individual predictions.

The present study demonstrates that item-level heterogeneity in neuroticism is reflected in distinct RSFC patterns, which align with both genetically defined clusters and psychometrically derived factor structures. These findings suggest that neuroticism is not a monolithic risk factor but comprises neurofunctionally and genetically distinct subcomponents that may differentially relate to psychological vulnerability.

From a clinical perspective, this insight provides a foundation for precision mental health approaches. For example, individuals exhibiting high scores on the Worry/Vulnerability or Anxiety/Tension factors may show greater propensity for specific cognitive-affective response styles (e.g., generalized anxiety, heightened vigilance, or social withdrawal), while those scoring higher on Depressed Affect-related connectivity patterns may be more likely to experience reduced motivation, slowed behavioral responses, or blunted emotional reactivity. Tailoring interventions to these subcomponents in both diagnostic assessment and treatment selection could improve specificity and effectiveness compared to conventional approaches that rely on composite trait scores or broad diagnostic categories.

In conclusion, RSFC association patterns revealed heterogeneous subcomponents of neuroticism that align with both genetic associations and psychometrically derived factor structures. These findings suggest that RSFC-based neurofunctional traits may serve as intermediate phenotypes, bridging heritable predispositions and self-reported psychological constructs. If RSFC reflects both relatively stable biological characteristics and state-dependent psychological processes, it could serve as a valuable biomarker for assessing both enduring traits and dynamic fluctuations in neuroticism. Future research should consider the dimensional and multifaceted nature of neuroticism and examine its associations with mental health outcomes at the facet-specific level, as different components may have distinct relationships with psychological sensitivity and adaptive functioning.

## Acknowledgments

This work was supported by the Laureate Institute for Brain Research. The funding sources had no role in the study design, analysis, decision to publish, or preparation of the manuscript. The authors are solely responsible for the content. During the preparation of this work the authors used ChatGPT (https://chat.openai.com/) and DeepL (https://www.deepl.com) in order to improve language and readability. After using these tools, the authors reviewed and edited the content as needed and take full responsibility for the content of the publication.

