## Supplementary for "Neuroticism Heterogeneity Through Item-Level Associations in Resting-State Functional Connectivity"

### Supplementary Materials for “Neuroticism Heterogeneity Through Item-Level Associations in Resting-State Functional Connectivity”

by Masaya Misaki, Chun Chieh Fan, Wesley K Thompson, Heekyeong Park, Bohan Xu, and Martin M Paulus

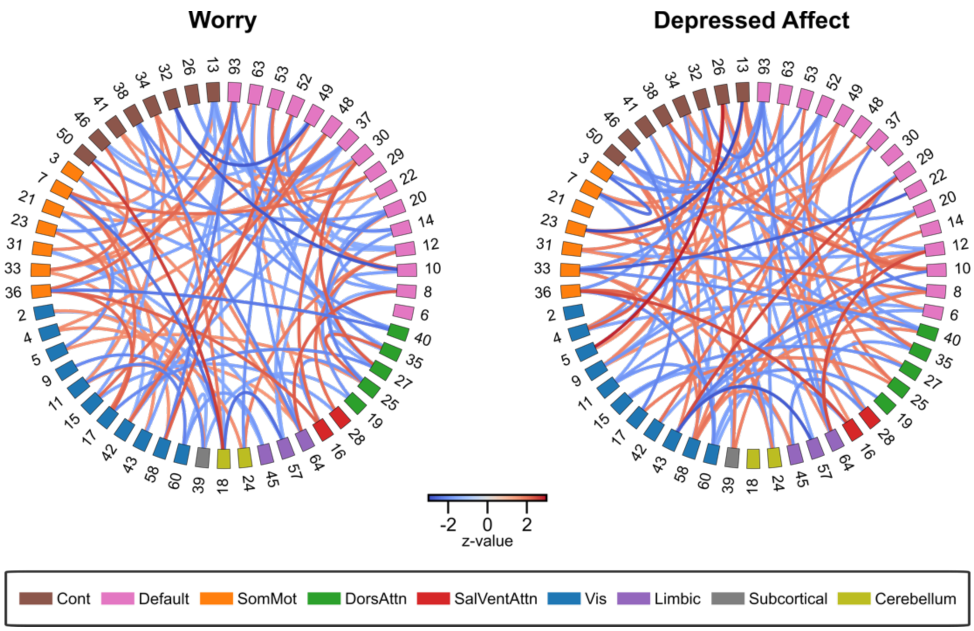


**Supplementary Figure S1**. Resting-state functional connectivity patterns associated with the Worry and Depressed Affect clusters of neuroticism items. The plot visualizes all connections with an absolute z-value greater than 0. Node numbers indicate the index of the independent component (IC), and node colors represent network labels. The IC weight maps are presented in Supplementary Figures S6A-S6I. The network labels were based on Yeo, Krienen (1): Cont, Frontoparietal Control network; Default, Default mode network; SomMot, Somatomotor network; DorsAttn, Dorsal attention network; Sal/VentAttn, Salience/Ventral attention network; Limbic, Limbic network; Vis, Visual network; Subcortical, Subcortical network; Cerebellum, Cerebellum network.

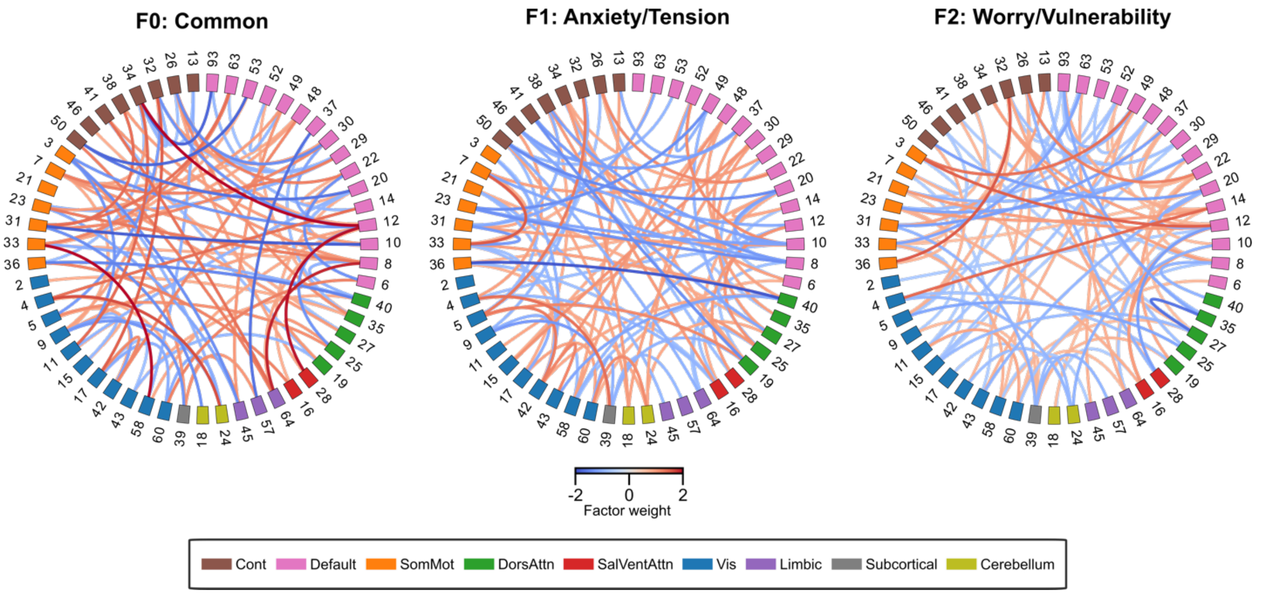


**Supplementary Figure S2**. Resting-state functional connectivity patterns associated with the factors of RSFC associations for neuroticism items, extracted through exploratory bi-factor analysis. The plot visualizes all connections with an absolute weighted z-value greater than 0. Node numbers indicate the index of the independent component (IC), and node colors represent network labels. The IC weight maps are presented in Supplementary Figures S6A-S6I. The network labels were based on Yeo, Krienen (1): Cont, Frontoparietal Control network; Default, Default mode network; SomMot, Somatomotor network; DorsAttn, Dorsal attention network; Sal/VentAttn, Salience/Ventral attention network; Limbic, Limbic network; Vis, Visual network; Subcortical, Subcortical network; Cerebellum, Cerebellum network.

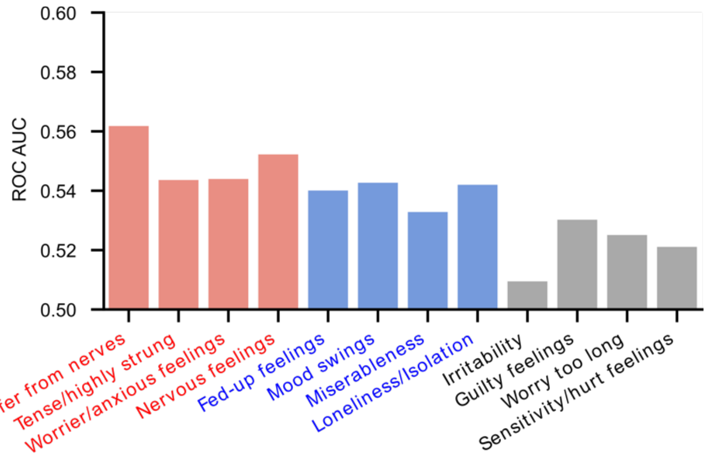


**Figure S3.** Generalization of the classification performance (Area Under the Receiver Operating Characteristic Curve; AUC) of the model trained on data from the first MRI instance to the second MRI instance. Colors indicate item clusters identified based on their genetic correlations [10]: Red represents the Worry cluster, while Blue represents the Depressed Affect cluster.

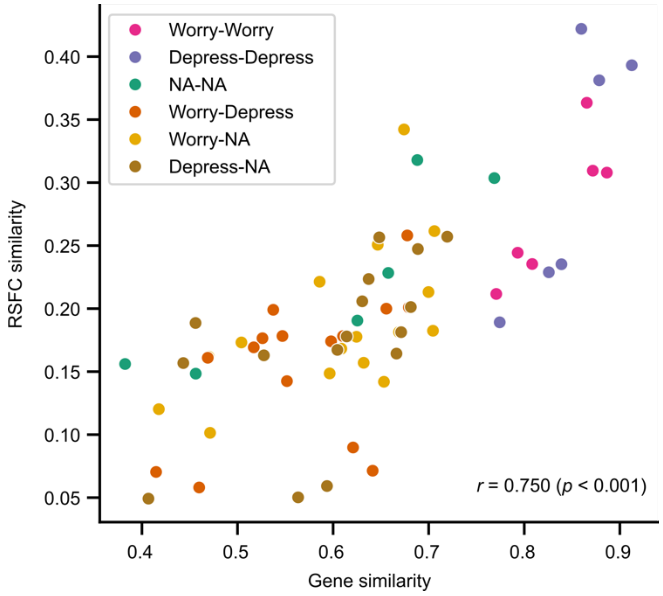


**Supplementary Figure S4**. The relationship between resting-state functional connectivity (RSFC) pattern correlations across items, derived from the second MRI visit data, and their genetic correlations obtained from Nagel, Watanabe (2). Dot colors indicate the combinations of item clusters, while NA denotes items that were not assigned to either cluster.

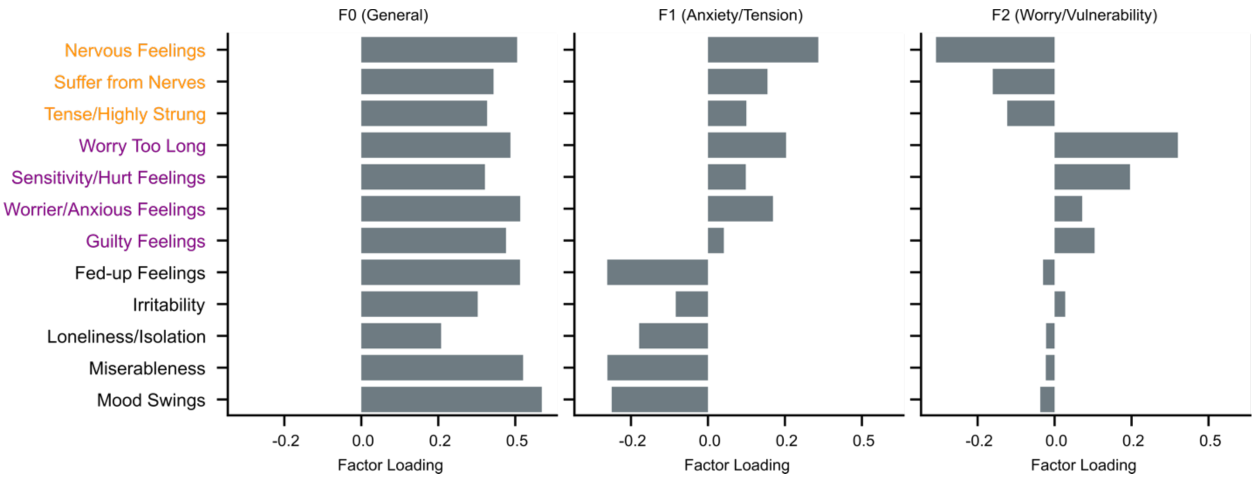


**Supplementary Figure S5**. Factor loadings for the general factor (F0) and two specific factors (F1, F2) extracted through exploratory bi-factor analysis of RSFC associations derived from the second MRI visit data. The colors of the row item labels indicate the specific factors identified based on questionnaire responses by Hill, Weiss (3): Orange represents the Anxiety/Tension factor, while Purple represents the Worry/Vulnerability factor.

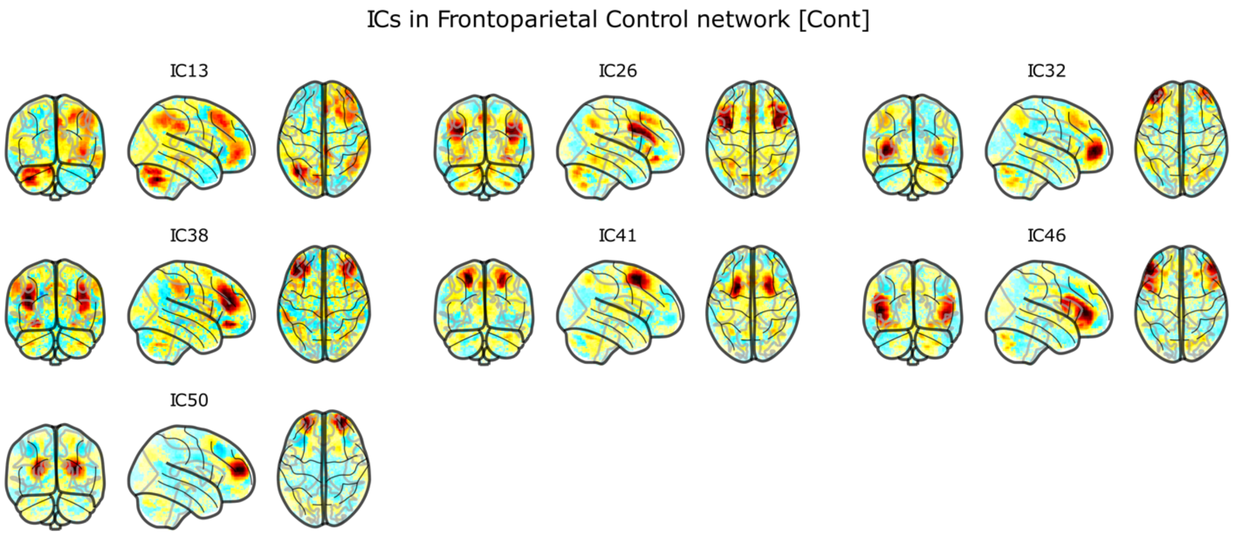


**Supplementary Figure S6a.** The independent component weight maps for the components in the Frontoparietal Control network shown on the glass brain.

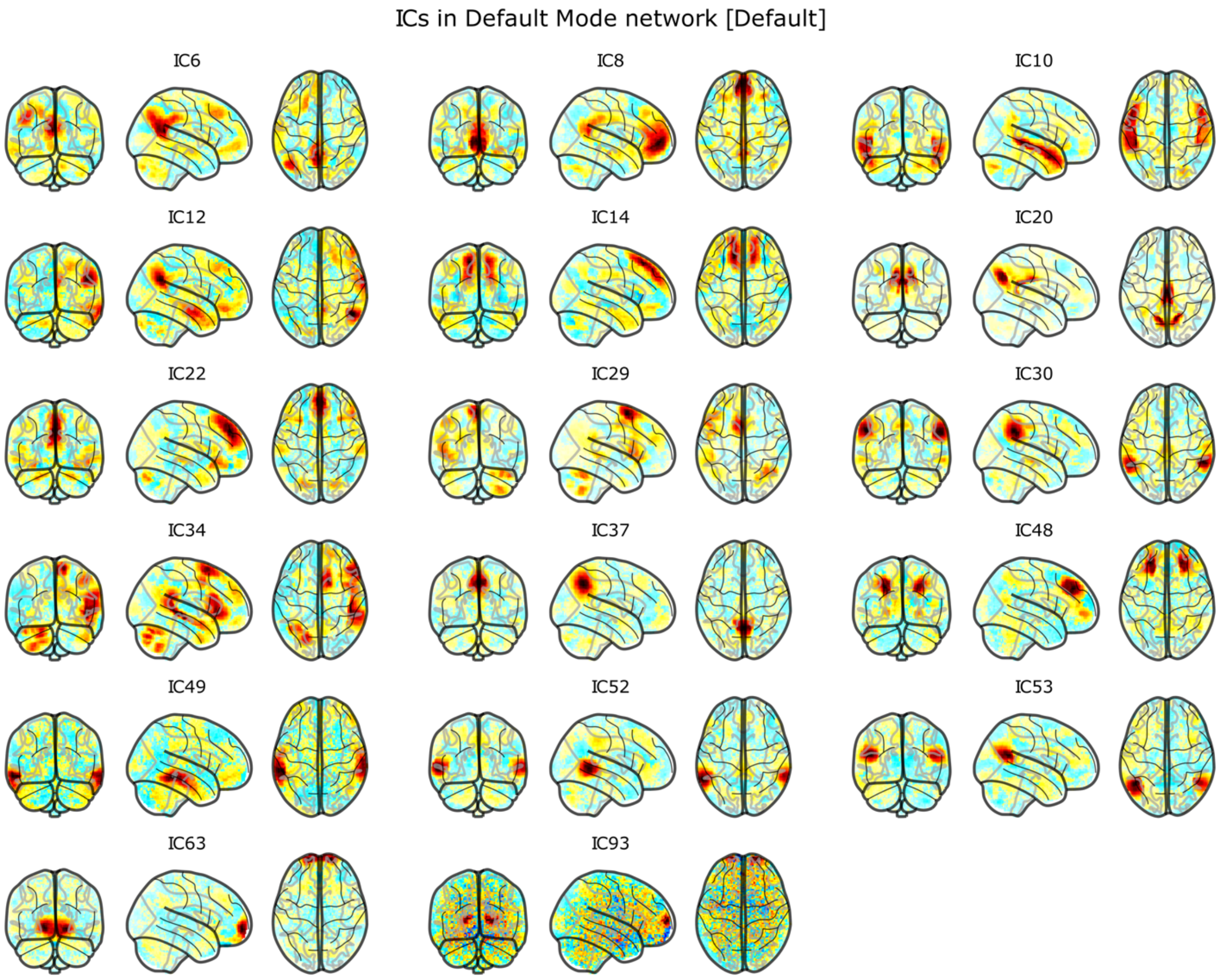


**Supplementary Figure S6b**. The independent component weight maps for the components in the Frontoparietal Control network shown on the glass brain.

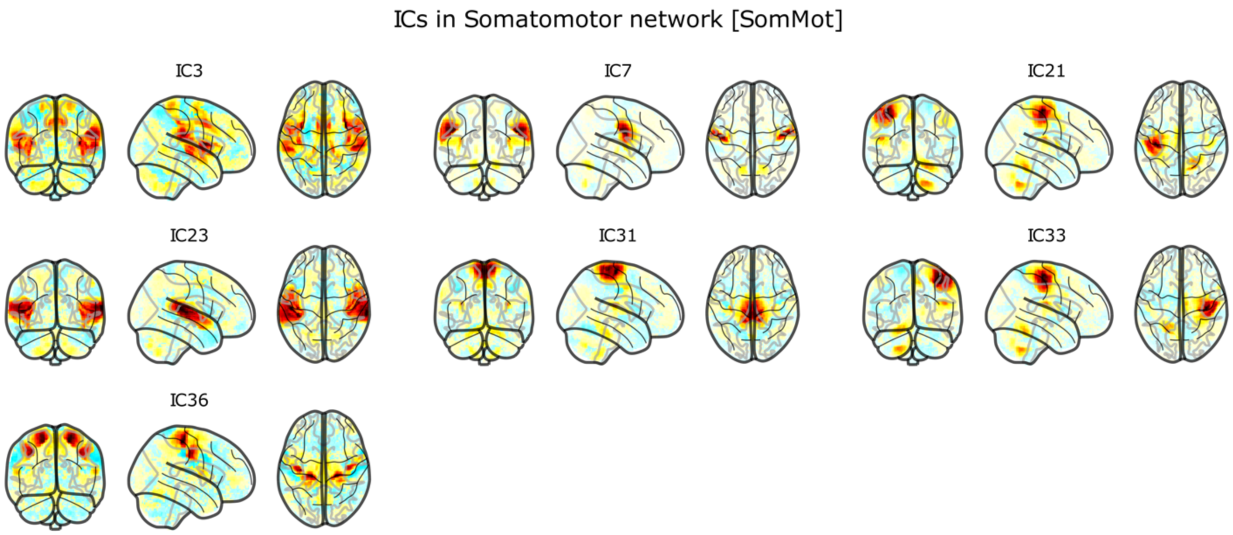


**Supplementary Figure S6c**. The independent component weight maps for the components in the Somatomotor network shown on the glass brain.

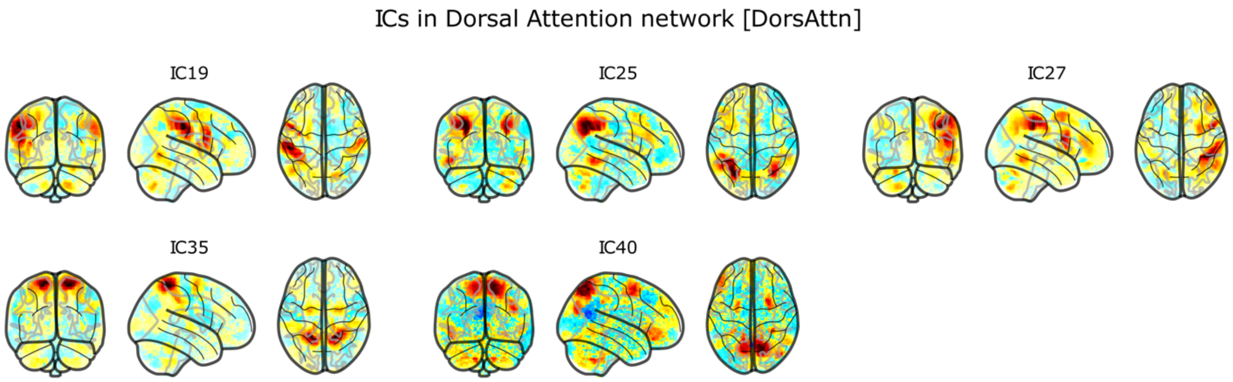


**Supplementary Figure S6D**. The independent component weight maps for the components in the Dorsal Attention network shown on the glass brain.

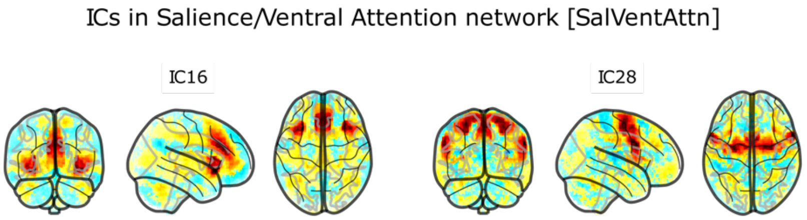


**Supplementary Figure S6e**. The independent component weight maps for the components in the Salience/Ventral Attention network shown on the glass brain.

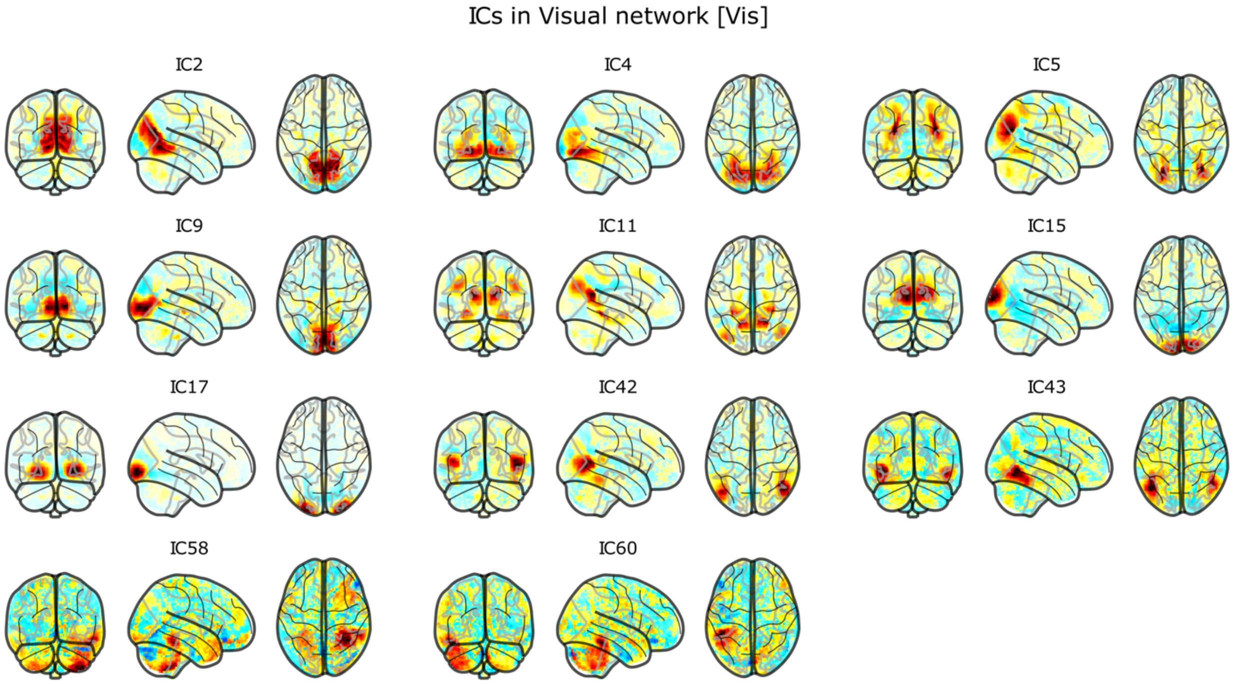


**Supplementary Figure S6f**. The independent component weight maps for the components in the Visual network shown on the glass brain.

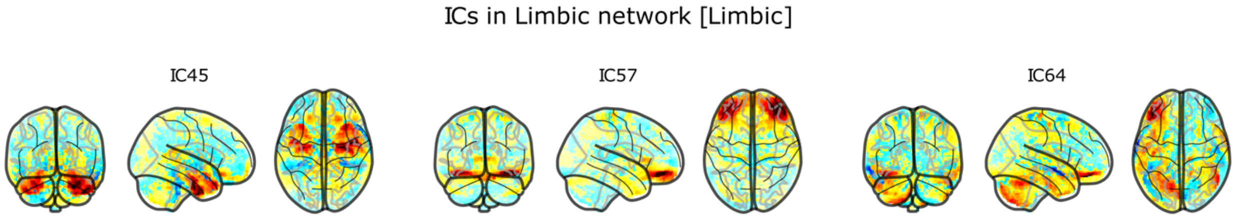


**Supplementary Figure S6g**. The independent component weight maps for the components in the Limbic network shown on the glass brain.

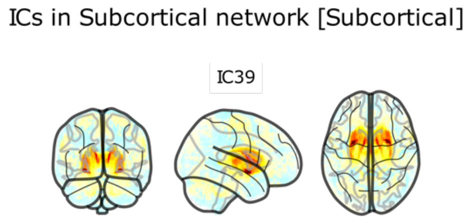


**Supplementary Figure S6h**. The independent component weight maps for the components in the Subcortical network shown on the glass brain.

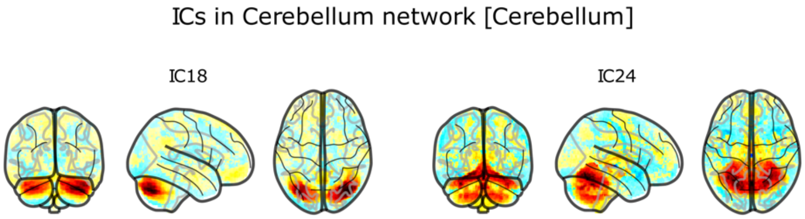


**Supplementary Figure S6i**. The independent component weight maps for the components in the Cerebellum network shown on the glass brain.
